# Pathogenicity, immunogenicity, and protective ability of an attenuated SARS-CoV-2 variant with a deletion at the S1/S2 junction of the spike protein

**DOI:** 10.1101/2020.08.24.264192

**Authors:** Pui Wang, Siu-Ying Lau, Shaofeng Deng, Pin Chen, Bobo Wing-Yee Mok, Anna Jinxia Zhang, Andrew Chak-Yiu Lee, Kwok-Hung Chan, Wenjun Song, Kelvin Kai-Wang To, Jasper Fuk-Woo Chan, Kwok-Yung Yuen, Honglin Chen

**Author notes:** **Correspondence to**: Honglin Chen, Department of Microbiology and State Key Laboratory for Emerging Infectious Diseases, Li Ka Shing Faculty of Medicine, The University of Hong Kong, Hong Kong SAR, China.

## Abstract

SARS-CoV-2 contains a PRRA polybasic cleavage motif considered critical for efficient infection and transmission in humans. We previously reported that virus variants with spike protein S1/S2 junction deletions spanning this motif are attenuated. Here we characterize a further cell-adapted SARS-CoV-2 variant, Ca-DelMut. Ca-DelMut replicates more efficiently than wild type or parental virus in cells, but causes no apparent disease in hamsters, despite replicating in respiratory tissues. Unlike wild type virus, Ca-DelMut does not induce proinflammatory cytokines in hamster infections, but still triggers a strong neutralizing antibody response. Ca-DelMut-immunized hamsters challenged with wild type SARS-CoV-2 are fully protected, demonstrating sterilizing immunity.

---

The emergence of SARS-CoV-2, a novel zoonotic origin β-coronavirus, has led to the first documented pandemic caused by a coronavirus ^1,2^. The virus continues to circulate globally in humans with rapidly increasing numbers of infections and casualties each day (https://coronavirus.jhu.edu/map.html). While coronaviruses from bats and pangolins have been found to be closely related to SARS-CoV-2 ^3-6^, the direct ancestral virus which attained cross-species transmission and the intermediate animal host source of human infections have not been defined. In the past two decades, three coronaviruses have jumped the species barrier to infect humans ^7^. SARS-CoV and MERS-CoV show limited human to human transmission ability while causing severe disease and mortality. In contrast, SARS-CoV-2, which uses the same human ACE2 binding receptor as SARS-CoV ^6^, is highly transmissible and causes variable severity of disease, from asymptomatic infections to severe and fatal outcomes. Analysis of the SARS-CoV-2 genome reveals a distinct PRRA polybasic cleavage motif at the S1/S2 junction of the spike protein when compared to the most proximal animal coronaviruses yet detected ^8^. Cleavage of spike into S1 and S2 subunits by the cellular protease furin promotes cell fusion mediated by S2, which is critical for cellular entry of coronavirus^9 10^. In avian influenza virus, acquisition of a polybasic cleavage motif facilitates infection of an increased variety of cell types ^11-13^. This polybasic cleavage site is essential for infection of human lung cells ^14^. It is speculated that acquisition of the PRRA polybasic motif in the spike protein of SARS-CoV-2 provides the virus with its unique ability of cross species transmission; this motif therefore appears to serve as a pathogenic element in human infections ^10,14^. If the PRRA polybasic cleavage motif was not a natural functional component in the original virus and acquired after cross species transmission, it is postulated that it may not be stable during the infection of its new hosts, or may need further adaptation. Indeed, a panel of SARS-CoV-2 variants with various lengths of deletion spanning the PRRA polybasic cleavage motif at the spike protein S1/S2 junction were identified in cultured cells and at low levels in clinical specimens, suggesting the S1/S2 junction may be under selection pressure as the SARS-CoV-2 virus circulates in humans ^15,16^. Initial characterization revealed that deletion at the S1/S2 junction causes SARS-CoV-2 virus attenuation in hamsters and prompted a further study to understand the properties of these deletion mutants in *in vitro* and *in vivo* systems. It remains to be determined whether some of these deletion variants may become more prevalent in humans as their circulation continues. On the other hand, these attenuated SARS-CoV-2 variants may hold promise as candidate live vaccines and it is important that their potential in this regard be evaluated.

To understand the role of the polybasic cleavage site in the infection and replication of SARS-CoV-2 and determine the stability of deletion variants, a previously characterized deletion mutant ^15^, Del-Mut-1, was further adapted in Vero cells to obtain a highly attenuated variant of SARS-CoV-2 virus, designated Ca-DelMut. We analyzed the growth properties of this variant in cells and evaluated its pathogenicity in a hamster model. While it is attenuated in the ability to cause disease in animals, Ca-DelMut replicates to a higher titer in Vero cells than the wild type SARS-CoV-2 virus. High titers of neutralizing antibodies were detected in animals previously infected with Ca-DelMut variant. However, infection with Ca-DelMut does not induce the high levels of proinflammatory cytokines seen in wild type virus infections. Importantly, Ca-DelMut immunized hamsters showed full protection with sterilizing immunity against challenge with two different strains of wild type virus.

## Results

### Growth and pathogenic properties of Ca-DelMut *in vitro* and *in vivo*

We previously identified and characterized a panel of SARS-CoV-2 variants containing 15-30bp deletions at the S1/S2 junction of the spike protein. One of the variants, Del-Mut-1, which contains a 30bp deletion spanningthe PRRA polybasic cleavage motif was shown to be attenuated in a hamster infection model ^15^. We further passaged this mutant in Vero E6 cells and obtained a variant with additional mutations in multiple genes in the background of Del-Mut-1, designated Ca-DelMut (**Figure 1 and Supplementary Table 1**). Growth properties of Ca-DelMut, parental Del-Mut-1 and a wild type (HK-13) SARS-CoV-2 were analyzed in Vero E6 and Calu-3 cells. It is interesting to note that in Vero E6 cells both Del-Mut-1 and Ca-DelMut grow to a significantly higher titer at the 24 and 48-hour time points than the wild type virus, with Ca-DelMut showing the strongest growth ability in both cell types (**Figure 2A**). Notably, Ca-DelMut exhibits a cold adaptation phenotype *in vitro*, as demonstrated by comparatively high level of replication at 30°C, whereas wild type virus (HK-13) replicates poorly at this temperature (**Figure S1**). Attenuation of Del-Mut-1 was reported previously ^15^. To further characterize the Ca-DelMut variant of SARS-CoV-2, we infected hamsters with variant and wild type SARS-CoV-2 strains. While inoculation with 10^3^ pfu HK-13 caused significant body weight loss post-infection, no apparent body weight loss was observed in Ca-DelMut (1.25×10^5^ and 10^3^ pfu) infected hamsters (**Figure 2B**). Histopathological analysis showed only mild regional alveolar septal infiltration and blood vessel congestion, with no obvious bronchiolar epithelium desquamation or luminal debris, no alveolar space infiltration or exudation, and no pathological changes in the intestines of Ca-DelMut infected hamsters, which are markedly different from the severe pathology observed in wild type virus infected animals (**Figure S2**). Examination of virus replication in lung and nasal turbinate tissues showed that Ca-DelMut replicates actively in the nasal turbinate tissues but that replication efficiency is much lower than that of HK-13 in hamster lungs (**Figure 2C**). These results indicate that Ca-DelMut has altered tissue tropism and is more likely to infect and replicate in the upper respiratory tract.

**Figure 1.**
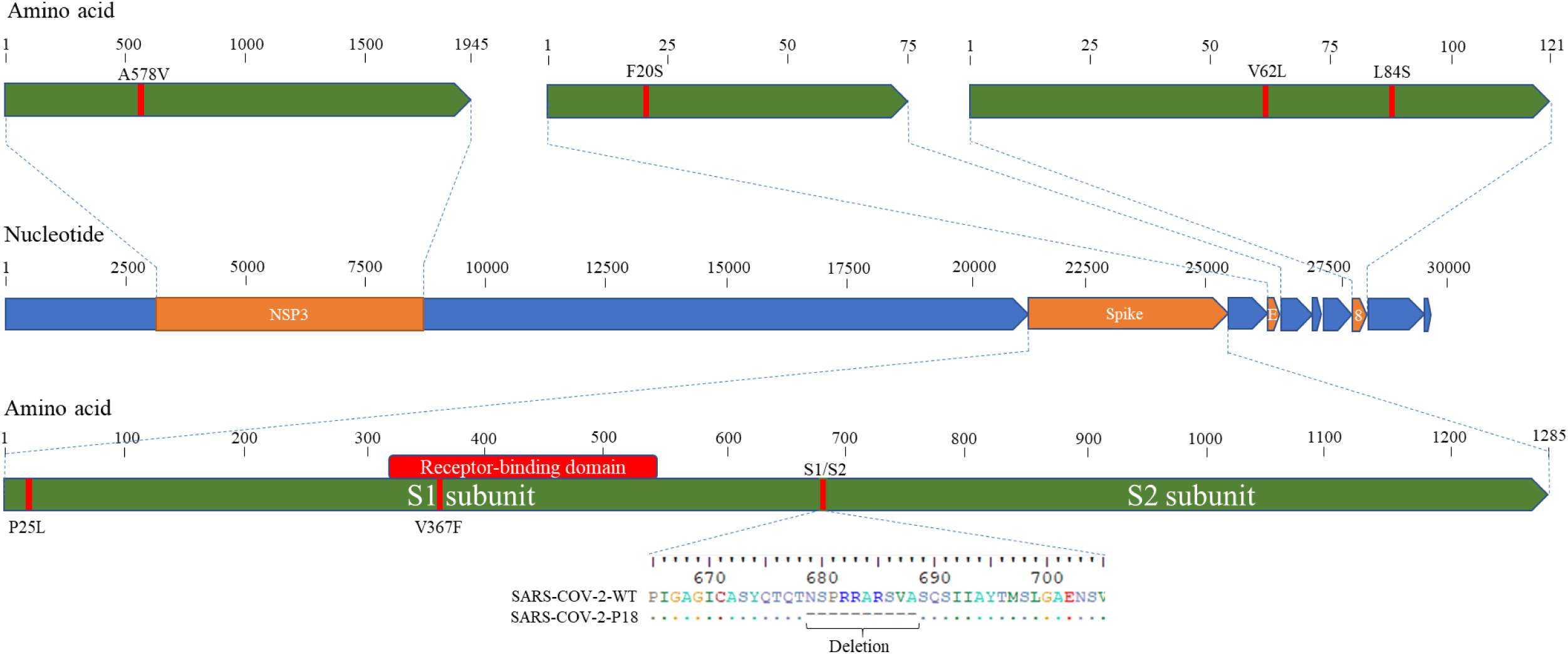
Schematic diagram of the SARS-CoV-2 genome showing the deletion and mutations of Ca-DelMut. Del-Mut-1 virus ^15^ was serially passaged in Vero E6 cells at 33°C (10 passages) and 30°C (8 passages). Virus from the 18^th^ passage was designated as Ca-DelMut live attenuated SARS-CoV-2 virus and amplified to prepare a virus stock. Ca-DelMut was sequenced by the Sanger method; mutations are shown in the diagram and in Supplementary Table 1.

**Figure 2.**
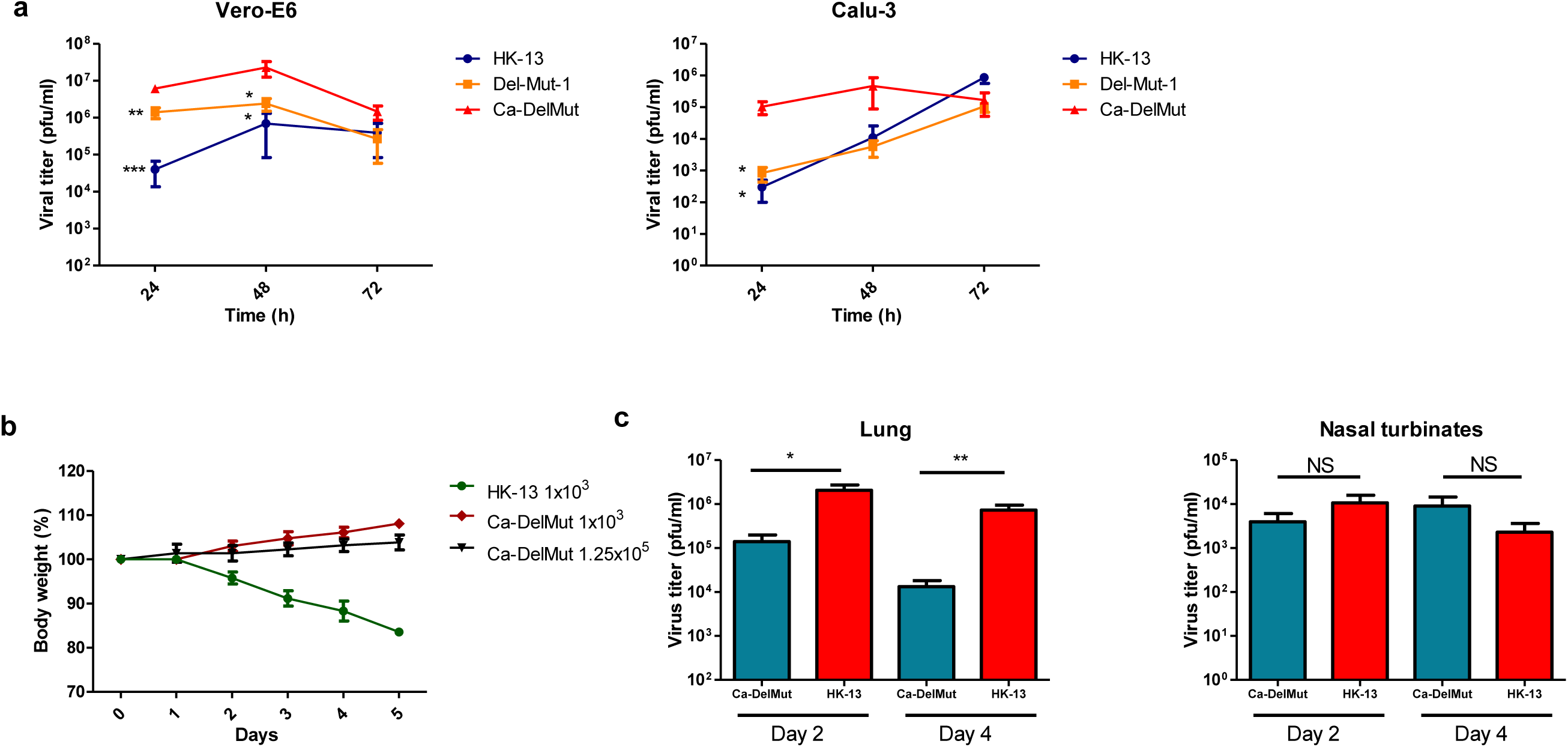
Replication efficiency of Ca-DelMut *in vitro* and *in vivo*. (A) Vero E6 or Calu-3 cells were infected with Ca-DelMut and other viruses at 0.01 moi and cultured at 37°C. At the indicated time points, supernatants were collected, and virus titer determined by plaque assay in Vero E6 cells. (B) Ca-DelMut infection in hamsters. Hamsters were infected intranasally with either Ca-DelMut or wild type (WT) viruses at different doses, as indicated. Body weight was monitored for 5 days. (C) Replication of Ca-DelMut in lung and nasal turbinate tissues. After virus challenge (1×10^3^ pfu), lung and nasal turbinate tissues were collected from hamsters at days 2 and 4, then homogenized and virus titer determined. Error bars represent mean ± s.d. (n=3). Statistical comparisons between means were performed by Student’s t-test: *** p<0.001, ** p<0.01, * p<0.05, NS: not significant.

### Immune response to infection with Ca-DelMut in hamsters

Impaired or dysfunctional immune responses have been characterized as an important mechanism of pathogenesis in human SARS-CoV-2 infections ^17-19^. Research on SARS-CoV and MERS-CoV has revealed that the interferon-mediated antiviral response is a double-edged sword which can induce both protective and pathogenic effects in humans. How Ca-DelMut infection induces the host immune response is of interest for understanding the molecular basis of its attenuation. To test if infection with Ca-DelMut may induce a different immune response from that elicited by wild type SARS-CoV-2, we examined interferon and cytokine expression in the lung tissues of virus infected hamsters. In contrast to infection with wild type HK-13 strain, we found that the Ca-DelMut variant does not provoke elevated levels of cytokines in infected hamsters (**Figure 3 and S3**). Aberrant activation of IL6 has been recognized as an important biomarker of disease severity in SARS-CoV-2 infected patients ^19-21^. Remarkably, activation of IL-6 was only observed in HK-13 strain infected hamsters but not in those infected with Ca-DelMut variant. We then analyzed the adaptive immune response in hamsters previously infected with Ca-DelMut or wild type virus. Levels of receptor binding domain (RBD) specific antibodies in sera collected three weeks after infection were determined. Sera from both hamsters previously infected with Ca-DelMut or that had recovered from wild type virus (HK-13) ^22^ infection showed robust induction of RBD specific antibodies (**Figure 4A and S4A**). Cell based neutralization assays also demonstrated strong neutralizing activity against the wild type virus strain HK-13 in sera collected from both Ca-DelMut variant and wild type SARS-CoV-2 virus (HK-13) infected hamsters (**Figure 4B and S4B**). These results indicate that infection with Ca-DelMut leads to an altered immune response which does not induce the elevated levels of proinflammatory cytokines seen in wild type SARS-CoV-2 virus infection. Nonetheless Ca-DelMut elicits a strong adaptive immune response, as demonstrated by robust neutralizing antibody production.

**Figure 3.**
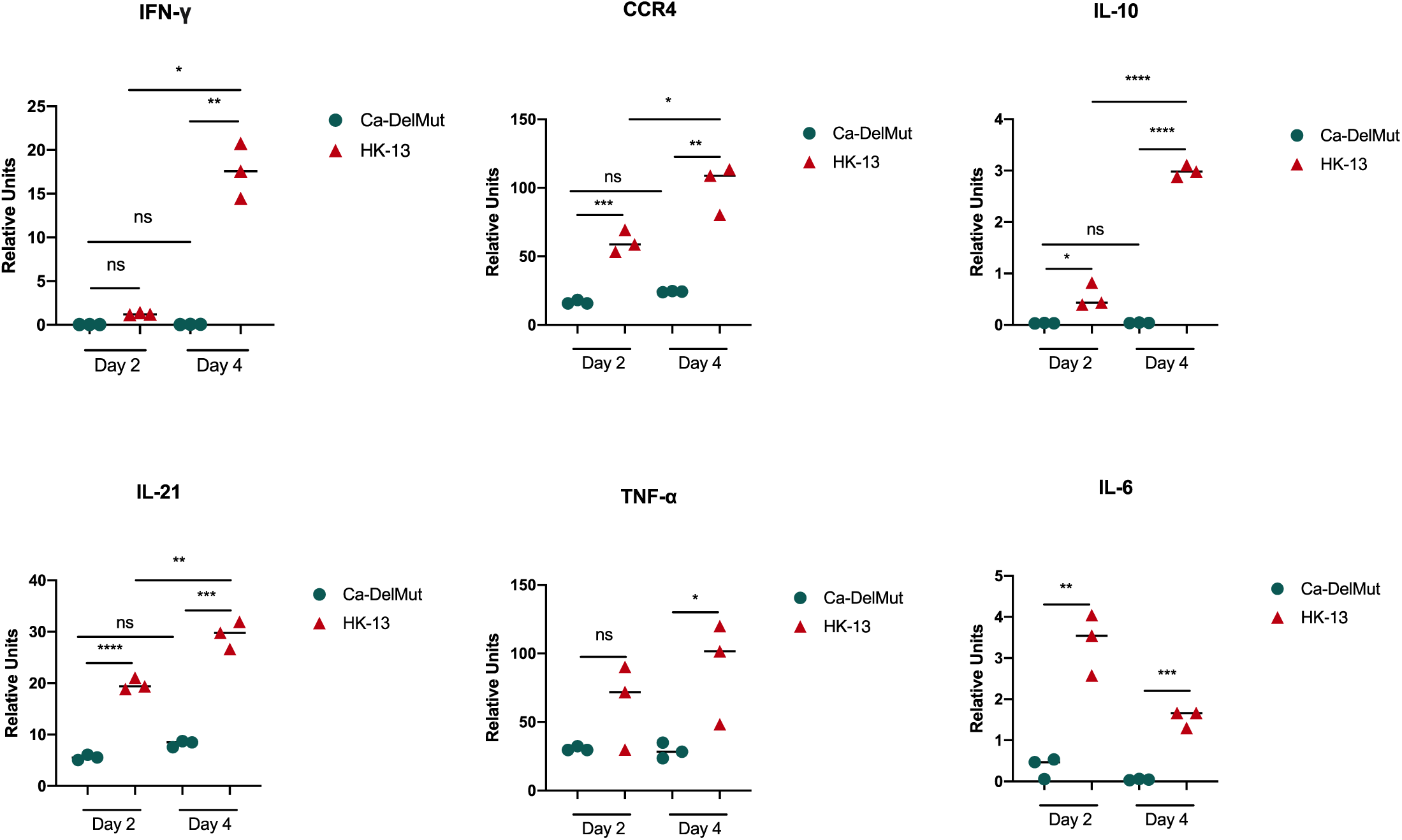
Proinflammatory cytokine response profiles in Ca-DelMut and wild type virus infected hamsters. Hamsters were infected intranasally with 1×10^3^ pfu of either Ca-DelMut or WT HK-13 virus. At days 2 and 4, RNA was extracted from lung tissues of infected hamsters and cDNA synthesized using oligo dT primers. Expression of different proinflammatory cytokines was examined by qPCR, normalized to an internal reference gene (hamster γ-actin), and the comparative Ct (2-ΔΔCt) method utilized to calculate the cytokine expression profile. Statistical comparisons between means were performed by Student’s t-test: **** p<0.0001, *** p<0.001, ** p<0.01, * p<0.05, ns: not significant.

**Figure 4.**
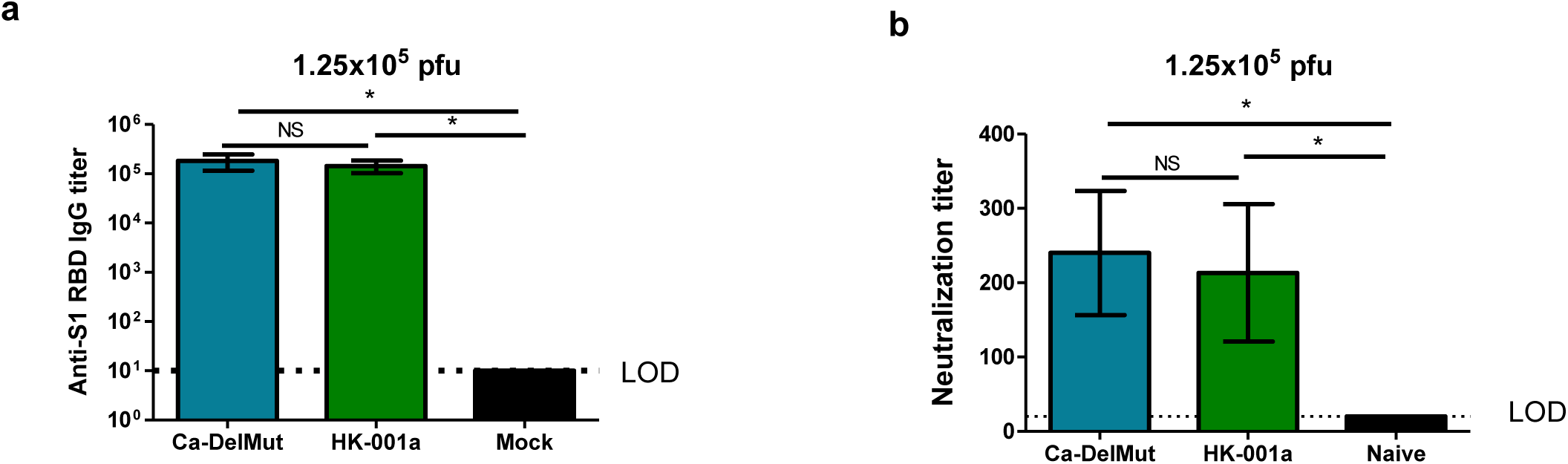
Antibodies induced by Ca-DelMut immunization of hamsters. Hamsters were immunized intranasally with 1.25×10^5^ pfu of either Ca-DelMut or wild type virus (HK-001a) virus^15,22^, or mock immunized. At day 21, blood was collected from hamsters and tested for anti-S1 RBD specific IgG titers and (B) neutralization activity against the HK-13 virus strain. Error bars represent mean ± s.d. (n=3). LOD: level of detection. Statistical comparisons between means were performed by Student’s t-test: * p<0.05, ns: not significant

### Infection with Ca-DelMut confers full protection against SARS-CoV-2 infection with sterilizing immunity

Because infection with Ca-DelMut causes no apparent disease in hamsters while inducing a strong neutralizing antibody response, we examined the potential of Ca-DelMut as a live attenuated virus vaccine to prevent SARS-CoV-2 virus infection and disease. Four weeks after infection with Ca-DelMut live attenuated virus, hamsters were re-challenged with wild type SARS-CoV-2 viruses. Two strains of SARS-CoV-2, HK-13 and HK-95, were used in the challenge experiment. HK-95 contains a D614G substitution in the spike protein, which has been suggested to bestow SARS-CoV-2 with higher infectivity in humans ^23^. Negligible body weight loss was observed in Ca-DelMut vaccinated hamsters infected with either strain of wild type virus, whereas control hamsters not immunized with Ca-DelMut lost about 12-15 percent of body weight by day 5 post-infection (**Figure 5A and S5**). Analyses of virus replication in the lung and nasal turbinate tissues of re-challenged hamsters showed that Ca-DelMut infection provides sterilizing immunity against subsequent challenge with either HK-13 or HK-95 SARS-CoV-2, with the quantity of virus being either very low or below the level of detection in both lung and nasal tissues on days 2 and 5 post-infection (**Figure 5B and S5**). Histopathological analysis showed that mock-vaccinated hamster lungs collected at day 5 post-infection with either wild type virus strain showed extensive alveolar exudation and infiltration; bronchiolar epithelial cell death with luminal exudation and cell debris were also observed (**Figure 6)**. In contrast, Ca-DelMut inoculated hamsters challenged with either strain of wild type SARS-CoV-2 virus only experienced mild regional alveolar septal infiltration and blood vessel congestion at days 2 and 5 post-infection. No other pulmonary histopathological changes were observed. Our experiment also showed a lower inoculum of Ca-DelMut (1 x 10^3^pfu) is sufficient to provide full protection to the wild type virus challenge (**Figure S6**). These data indicate that prior infection with live attenuated Ca-DelMut variant virus provides complete protection against infection with wild type SARS-CoV-2 viruses in hamsters.

**Figure 5.**
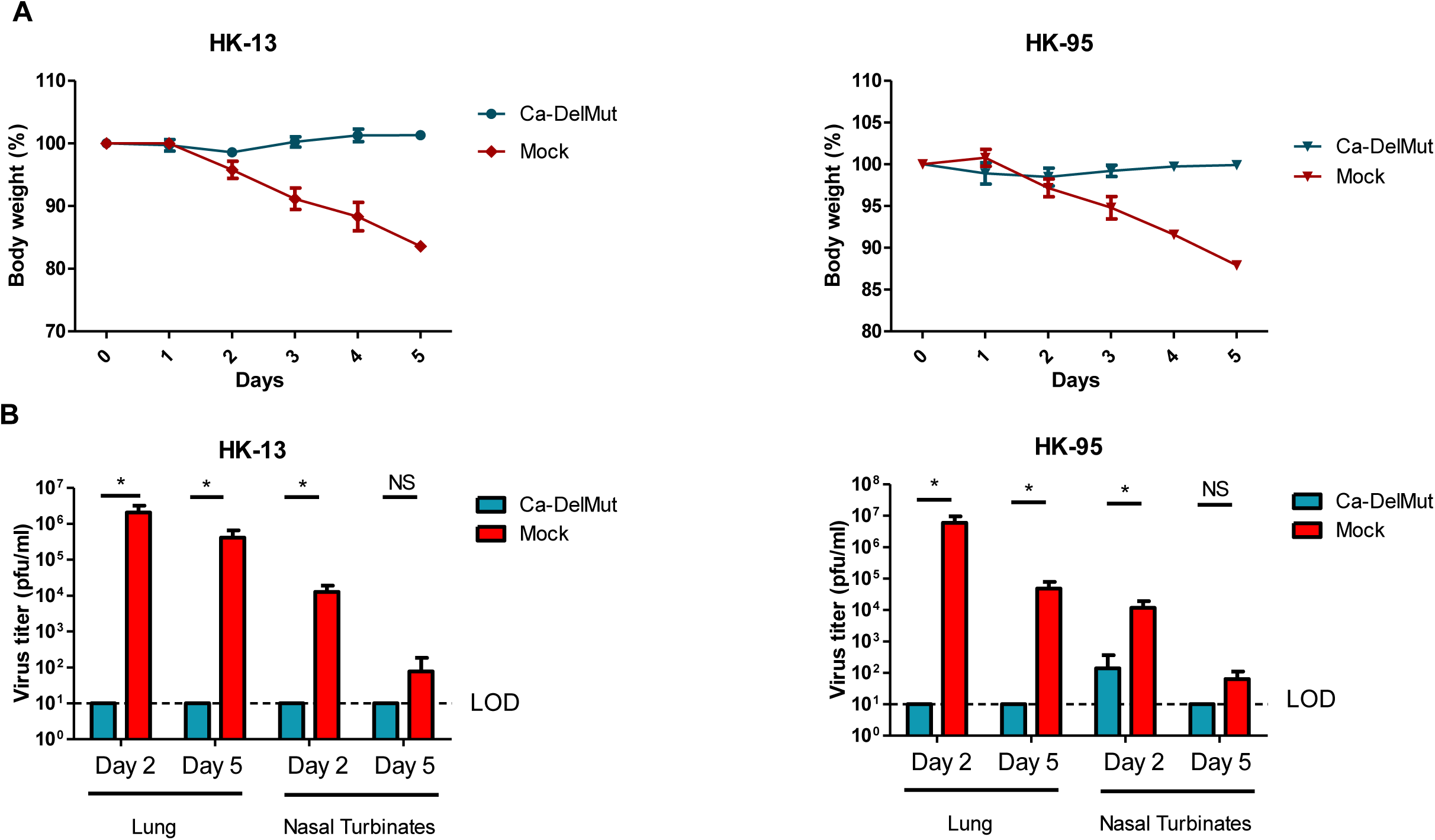
Ca-DelMut immunization protection against WT virus challenge in hamsters. Hamsters were inoculated with 1.25×10^5^ pfu Ca-DelMut or mock immunized. At day 28 after immunization, hamsters were challenged with 1×10^3^ pfu of either HK-13 or HK-95 virus. (A) Body weight and disease symptoms were monitored for 5 days. (B) At days 2 and 5 post-infection, lungs and nasal turbinate tissues were collected for virus titration and histopathological study. Error bars represent mean ± s.d. (n=3).

**Figure 6.**
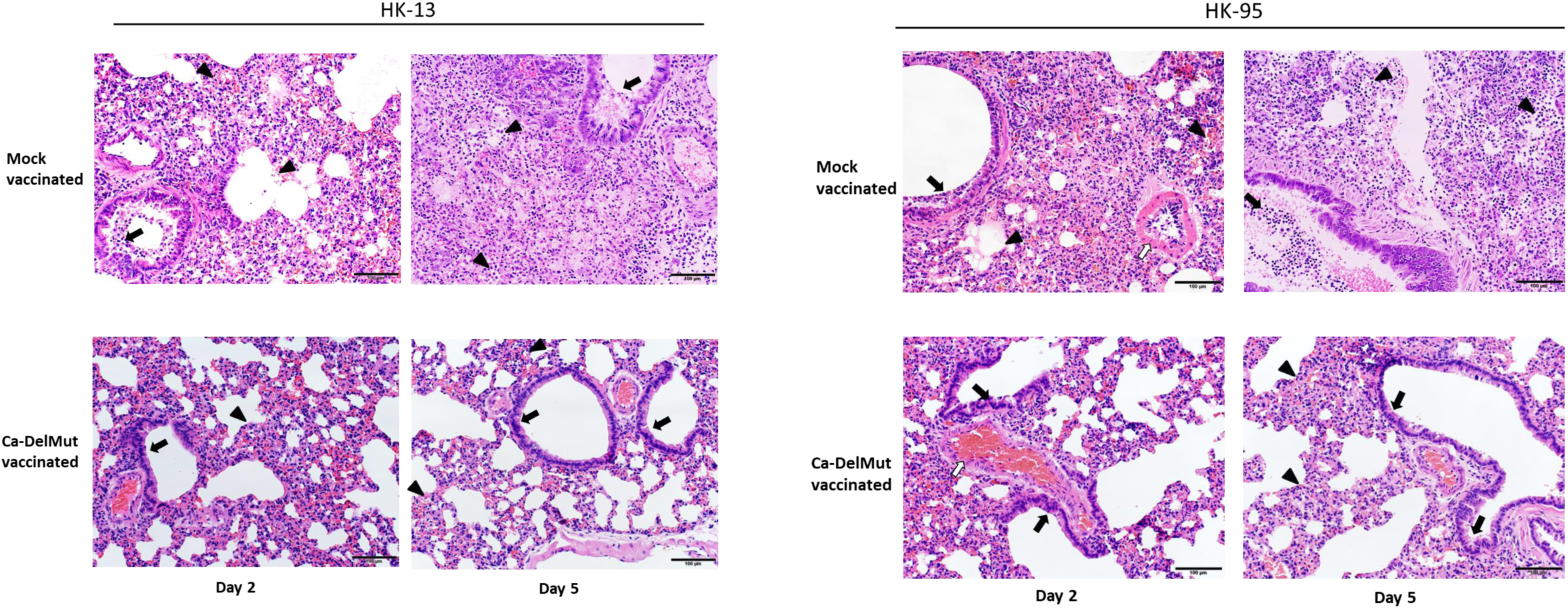
Histopathological analysis of lung pathology in WT virus challenged Ca-DelMut-and mock-immunized hamsters. At day 28 after immunization, hamsters were challenged with 1×10^3^ pfu of either HK-13 or HK-95 virus. At days 2 and 5 post-infection, lungs were collected, fixed, processed into paraffin blocks and sections H&E stained. Challenge with HK-13. Day 2: Mock-vaccinated hamster lungs showed bronchiolar epithelial cell death and the bronchiolar lumen filled with exudate and cell debris (arrow). Diffuse alveolar infiltration and focal hemorrhage were also seen (arrowheads). Ca-DelMut-vaccinated hamster lungs showed regional alveolar septal infiltration and blood vessel congestion (arrowhead), but no obvious bronchiolar epithelial cell death (arrow). Day 5: Lungs of mock-vaccinated hamsters showed severe alveolar infiltration and exudation (arrowheads), as well as bronchiolar luminal exudation (arrow). Vaccinated hamster lungs showed focal alveolar septal infiltration (arrowheads), while the bronchiolar epithelium appeared normal with no luminal secretion or cell debris (arrows). (b)Challenge with HK-95. Day 2: Mock-vaccinated hamster lungs showed bronchiolar epithelial cell death with luminal cell debris (arrow) and diffuse alveolar infiltration with focal hemorrhage and exudation (arrowheads), with a medium sized blood vessel showing severe endotheliitis (open arrow). Vaccinated hamster lungs showed regional alveolar septal infiltration and blood vessel congestion (arrowhead); a blood vessel appeared to be normal (open arrow), and no obvious bronchiolar epithelial cell death was observed (arrows). Day 5: Lungs of mock-vaccination control hamsters showed severe alveolar infiltration and exudation (arrowheads) and bronchiolar luminal cell debris (arrow). Vaccinated hamster lungs showed focal alveolar septal infiltration (arrowheads), while the bronchiolar epithelium appeared normal without luminal secretion (arrows). Scale bar: 100μm.

## Discussion

Coronaviruses are zoonotic pathogens with distinct cross species transmissibility ^24^. Besides 229E, OC43, HKU-1 and NL63, which have long been circulating in humans, SARS-CoV, MERS-CoV and SARS-CoV-2 have jumped the species barrier to infect humans in recent years ^7^. SARS-CoV disappeared in 2004, while MERS-CoV is restricted to a few countries in the Middle East with only sporadic human transmissions since 2012 ^25^. SARS-CoV-2 virus utilizes the same cellular receptor as SARS-CoV, ACE2, for mediating human infection but has exhibited a distinctive infectivity and transmissibility profile since it was first recognized in humans in Wuhan, China, in December 2019 ^6^. There is strong interest in understanding how SARS-CoV-2 has acquired the unique ability to infect and transmit efficiently in humans. SARS-CoV-2 contains a PRRA polybasic motif not seen in the most closely related bat and pangolin coronaviruses currently known ^3,5,8^. The presence of a polybasic cleavage site at the S1/S2 junction of the spike protein of SARS-CoV-2 virus is considered a critical property for enhanced coronavirus infectivity in humans and zoonotic potential ^14,26^. A peptide assay has shown that the polybasic cleavage site harbored in SARS-CoV-2 is more accessible to proteases which activate the coronavirus spike protein ^10^. If the polybasic cleavage site is one of the essential elements providing SARS-CoV-2 with increased infectivity and pathogenicity in humans, removal of this determinant could logically attenuate SARS-CoV-2 into a mild respiratory virus similar to the less pathogenic common coronaviruses currently circulating. Our previous report on a panel of attenuated variants with deletions at the S1/S2 junction supports this contention ^15^. This study has further characterized one such mutant, Ca-DelMut, showing that it has low pathogenicity and does not provoke an inflammatory response in hamsters and that it induces adaptative immunity protective against subsequent infection with more pathogenic SARS-CoV-2 strains. Ca-DelMut attenuated virus could also be a useful tool for studying SARS-CoV-2 replication, host tissue tropism and transmissibility.

Although both SARS-CoV and SARS-CoV-2 utilize ACE2 to mediate infection, the distinctive infectivity and pathogenicity displayed by SARS-CoV-2 is likely to be associated with the acquisition of a polybasic furin cleavage site in the protein which, together with enhanced binding affinity of the SARS-CoV-2 RBD for ACE2, would significantly broaden the tissue tropism of this virus^14,27^. Deregulated innate immunity during the early stage of infection in the upper respiratory tract may determine the subsequent outcome of dissemination to the lower respiratory tract and disease severity ^17,28^. We found that Ca-DelMut replicates to comparable levels to wild type virus in the nasal turbinates but less effectively than wild type virus in the lungs (**Figure 2C**). Importantly, while replication of Ca-DelMut variant was observed in lung tissues, the elevated expression of proinflammatory cytokines elicited in wild type virus infected hamsters was not detected (**Figure 3 and S3**). These observations clearly suggest that the polybasic cleavage motif is a virulence element in SARS-CoV-2 and that its removal makes SARS-CoV-2 much less pathogenic, and more similar to a common cold respiratory coronavirus. We believe that SARS-CoV-2 will continue to undergo further adaptation as it circulates in humans. Our previous study revealed that variants with deletions at the S1/S2 junction are present at low levels in clinical specimens ^16^. In 2003, the SARS-like coronavirus characterized from civet cats and early-outbreak human SARS-CoV isolates contained a 29-bp sequence in the ORF8 sequence which was deleted following its subsequent circulation in humans ^29^. Deletions in ORF7b and ORF8 have also been observed in SARS-CoV-2, although the significance of these alterations is currently unknown ^30,31^. It remains to be seen if continued evolution of SARS-CoV-2 in humans will subsequently select for less pathogenic variants similar to Ca-DelMut, or with other mutations.

Several SARS-CoV-2 vaccines are being developed for use in humans ^32-35^. However, no vaccine against a coronavirus has been developed before and there are concerns regarding whether current vaccine strategies will be able to provide adequate and sufficiently long lasting immunity to prevent infection and alleviate disease severity and spread. Anti-SARS-CoV-2 antibodies are reported to decline rapidly in naturally infected individuals ^36,37^. This study showed that Ca-DelMut induces a different innate immune response to that observed in wild type virus infections in an animal model and evokes sterilizing protective adaptative immunity against challenge with wild type virus (Figure 4). Because Ca-DelMut is attenuated it does not provoke proinflammatory cytokines which could interfere with the induction of adaptive immunity. It is possible that the Ca-DelMut variant may be able to stimulate more balanced and long-lasting immunity; further study is required to comprehensively compare the immune profiles induced by Ca-DelMut and wild type SARS-CoV-2 viruses. The potential applications of this attenuated SARS-CoV-2 virus should be explored and evaluated. However, given the high replication efficiency and low pathogenicity of Ca-DelMut, it may be an ideal strain for production of an inactivated vaccine, in addition to holding promise as a live attenuated vaccine.

## Online Methods

### Generation of cell adapted Del-mut (Ca-DelMut) virus

Del-mut viruses were prepared as previously described ^15^. To generate Ca-DelMut virus, the Del-Mut-1 virus was serially passaged 10 times in Vero E6 cells at 33°C and then passaged at 30°C 8 times. For each passage, incubation time was 2-3 days, depending on the occurrence of cytopathic effects. After the 18^th^ passage, the virus was amplified in a T75 Flask, then titered by plaque assay and sequenced using the Sanger method.

### Growth kinetics

Confluent Vero E6 or Calu-3 cells were infected at 0.01 moi with the indicated viruses and incubated for 3 days. Virus supernatant was collected at the indicated time points. Virus titers were determined by plaque assay using Vero E6 cells.

### Plaque assay

Confluent Vero E6 cells in 6-well format were incubated with 10-fold serially diluted virus for 1 h. After adsorption, virus was discarded. Cells were washed and overlaid with 1% agarose in DMEM and incubated for 3 days at 37°C. Cells were fixed with 10% formaldehyde for 1 day. Agarose gels were then removed and plaques stained with 1% crystal violet and counted.

### Immunization and challenge of hamsters

7-8 week old golden Syrian hamsters were anesthetized intraperitoneally with ketamine and xylazine and then immunized intranasally with 1.25×10^5^ pfu of Ca-DelMut or SZ-002 virus or mock immunized with PBS. Body weight and disease symptoms were monitored daily. At day 21, sera were collected from hamsters for anti-spike RBD IgG and neutralizing antibody determination. At day 28, Ca-DelMut- and mock-immunized hamsters were challenged with WT virus (HK-13 or HK-95) at a dose of 1×10^3^ pfu. Body weight and disease symptoms were monitored daily and at days 2 and 5, lung and nasal turbinate tissues were collected for histopathology and virus titration by plaque assay. For low dose immunization, hamsters were immunized with 10^3^ pfu of either Ca-DelMut or WT HK-13 virus, or mock immunized with PBS. All experiments involving SARS-CoV-2 were conducted in a biosafety level 3 laboratory. All animal studies were approved by the Committee on the Use of Live Animals in Teaching and Research, The University of Hong Kong.

### Pathogenicity of Ca-DelMut and wild type SARS-CoV-2 virus in hamsters

Hamsters were challenged intranasally with 1×10^3^ or 1.25×10^5^ pfu of Ca-DelMut or 1×10^3^ pfu of WT virus. Body weight and disease symptoms were monitored daily. At days 2 and 4, lung and nasal turbinate tissues were collected for histopathological study, determination of proinflammatory cytokine expression and virus titration.

### Neutralization assay

Heat inactivated sera from Ca-DelMut challenged hamsters were 2-fold serially diluted in DMEM medium and incubated with 100 pfu of the indicated virus at 37°C for 1 h. The mix was added to confluent Vero E6 cells and incubated for 4 days at 37°C. Naïve and WT challenged sera were used as controls. After 4 days, cytopathic effect (CPE) was detected by microscopy, with the neutralization endpoint being the highest serum dilution causing 50% inhibition of CPE.

### ELISA

A hamster anti-spike RBD IgG detection kit (Wantai-Bio) was used to detect RBD specific antibodies. Procedures were conducted in accordance with the manual. Briefly, heat inactivated sera from Ca-DelMut challenged hamsters were 10-fold serially diluted and added to the plate and incubated at 37°C for 30 mins. Sera from mock- and WT-challenged hamsters were included as controls. The plate was washed 5 times and then incubated with secondary antibody reagent at 37°C for 30 mins. After washing, color development solution was added and the plate incubated at 37°C for 15 mins. Stop solution was added and absorbance at 450 nm measured.

### Quantification of expression of proinflammatory cytokines and chemokines

Expression of proinflammatory cytokines was quantified using a qRT-PCR technique similar to that described in a previous study ^38^. Briefly, total RNA was extracted from hamster lungs using RNAzol RT reagent (MRC) according to the manual. cDNA was synthesized using a High Capacity cDNA Reverse Transcription Kit (Invitrogen) and oligo dT primers following the protocol provided. qPCR was performed using SYBR Premix Ex Taq (Takara) reagent and gene specific primers in an LC480 PCR machine (Roche). PCR conditions were as follows: initial denaturation: 95°C for 5 min, 45 cycles of amplification: 95°C for 10s, 60°C for 10s, 72°C for 10s, and melting curve analysis: 65°C to 97°C at 0.1°C/s. Expression of target genes was normalized to the internal reference gene (hamster γ-actin) and the comparative Ct (2-ΔΔCt) method utilized to calculate the cytokine expression profile.

### Histopathology

Organs were fixed in 10% PBS buffered formalin and processed into paraffin-embedded blocks. Tissue sections were stained with haematoxylin and eosin (H&E) and examined by light microscopy as in a previous study^38^.

## Acknowledgements

The authors would like to thank Dr Jane Rayner for critical reading and editing of the manuscript. This study is partly supported by the Theme-Based Research Scheme (T11/707/15) and General Research Fund (17107019) of the Research Grants Council, Hong Kong Special Administrative Region and the Sanming Project of Medicine in Shenzhen, China (No. SZSM201911014).

## Author Contributions

P.W., and H.C. designed the studies, P.W., S-Y. L., S. D., P. C., B.W.M., A.J.Z., A.C-Y.L.,K-H.C. and W.S. performance experiments; P.W., S-Y. L., A. J. Z., K. K-W.T., J. F-W.C, K-Y. Y. and H.C. analyzed the data; P.W. and H.C. wrote the paper.

---

**Supplementary Table 1.** Deletion and mutations in Ca-DelMut virus genome compared to Wuhan-Hu-1

| Nucleotide coordinate | Wuhan-Hu-1 | Ca-DelMut |  |  |  |
| --- | --- | --- | --- | --- | --- |
|  |  | Nucleotide | Amino acid | Protein | ORF |
| 1663 | C | T | No change | NSP2 | ORF1a |
| 4455 | C | T | A578V | NSP3a | ORF1a |
| 8782 | C | T | No change | NSP4 | ORF1a |
| 21636 | C | T | P25L | S | S |
| 22661 | G | T | V367F | S | S |
| 23598-23627 | No deletion | 30-base-pair deletion | 10 aa deletion | S | S |
| 24034 | C | T | No change | S | S |
| 26303 | T | C | F20S | E | E |
| 26729 | T | C | No change | M | M |
| 28077 | G | C | V62L | ORF8 | ORF8 |
| 28144 | T | C | L84S | ORF8 | ORF8 |

\*Details of deletion and mutations in the Ca-DelMut variant with reference to the Wuhan-Hu-1 SARS-CoV-2 strain (Wu et al., 2020)
Wu, F., Zhao, S., Yu, B., Chen, Y.M., Wang, W., Song, Z.G., Hu, Y., Tao, Z.W., Tian, J.H., Pei, Y.Y., *et al.* (2020). A new coronavirus associated with human respiratory disease in China. *Nature* 579, 265-269.

**Figure S1.**
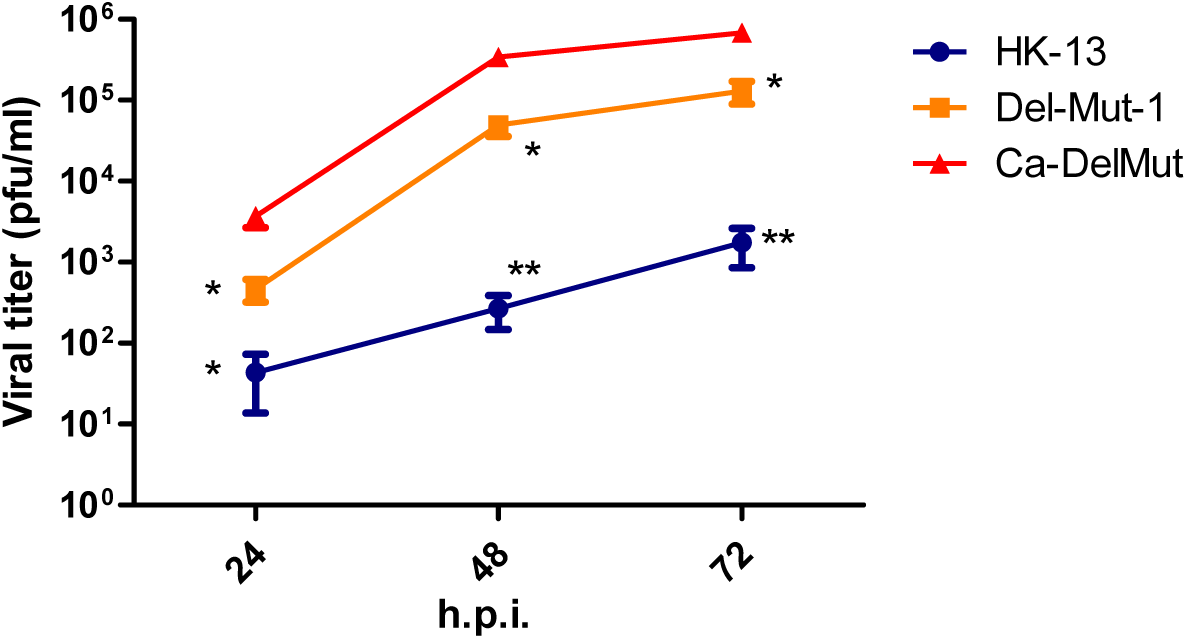
Ca-DelMut exhibits a cold-adapted phenotype *in vitro*. Vero E6 cells were infected with Ca-DelMut and other viruses at 0.01 moi and incubated at 30°C. At the indicated time points, supernatants were collected and virus titer determined by plaque assay. Error bars represent mean ± s.d. (n=3). h.p.i.: hours post infection.

**Figure S2.**
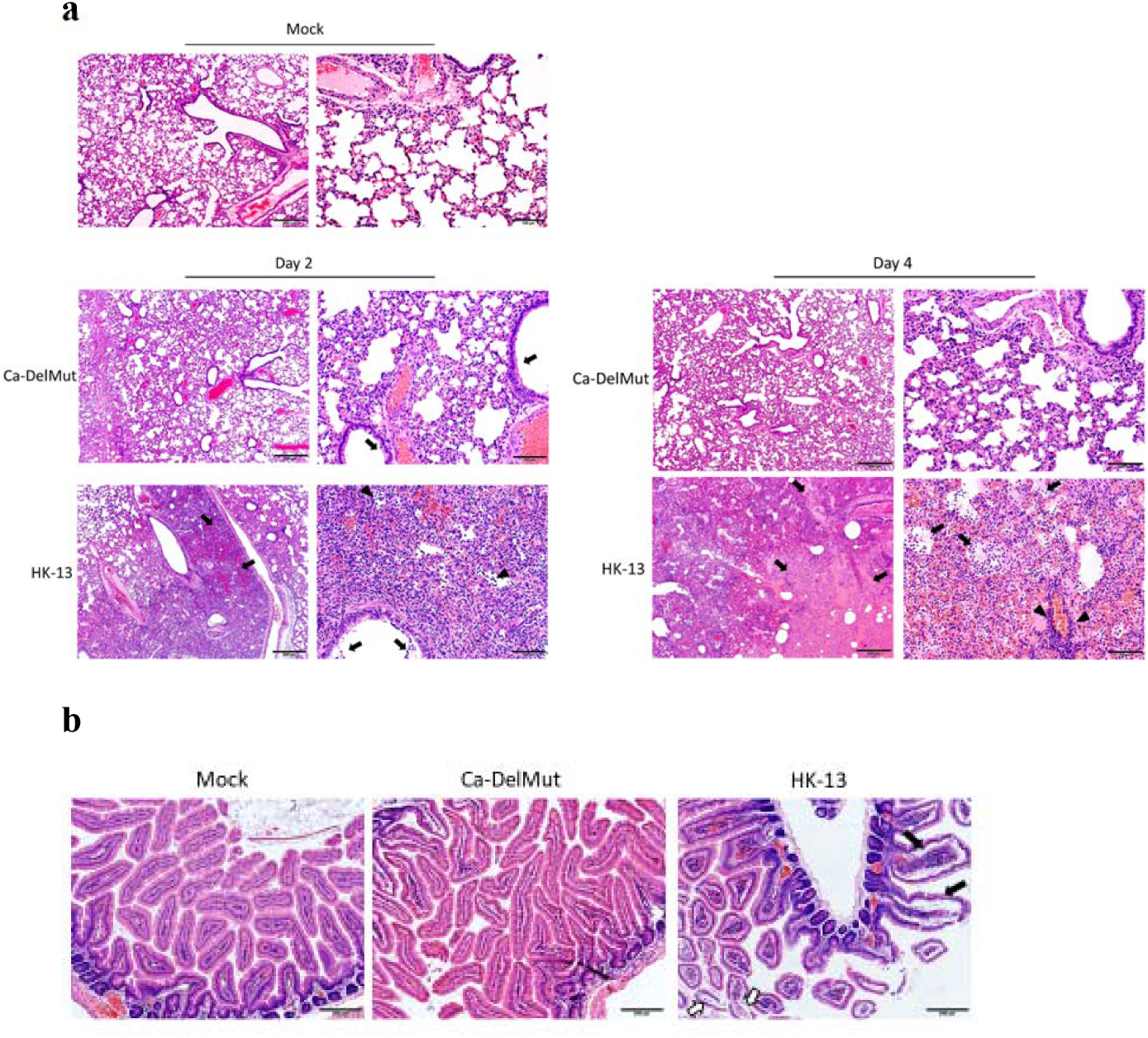
Infection with Ca-DelMut causes only mild pathological changes. Hamsters were infected with 1×10^3^ pfu of either Ca-DelMut or HK-13 (WT), or mock infected. At days 2 and 4 post-infection, lungs and small intestine were collected and fixed in 10% formalin, and then processed into paraffin blocks and sections H&E stained. (a)The top panel shows normal lung structures in the lungs of mock-infected control hamsters at 4x (left) and 20x (right) magnification. At day 2 after infection with Ca-DelMut, lung tissues showed only mild regional alveolar septal infiltration and blood vessel congestion. No obvious bronchiolar epithelium desquamation or luminal debris (arrows), and no alveolar space infiltration or exudation were observed. At day 4, no deleterious progression of histopathology was observed. For WT virus at day 2 post-infection, the low magnification image (left) showed regional lung consolidation and focal pulmonary hemorrhage (arrows). The higher magnification image (right) showed massive alveolar space infiltration (arrowheads) and hemorrhage, with a little bronchiolar luminal cell debris (arrows) visible. At day 4 post WT**-**infection, the low magnification image (left) showed intensive alveolar exudation, infiltration and hemorrhage resulting in pulmonary consolidation (arrows), while the higher magnification image (right) showed intensive protein rich exudates filling the alveolar space (arrows), in addition to massive infiltration and alveolar hemorrhage; a blood vessel shows moderate infiltration (arrowheads). (b)Representative histological images of hamster small intestines. The left-most image shows mock-infected control hamster small intestinal villi with normal structure. At day 4 post infection with Ca-DelMut, no apparent histopathological changes were detected in the small intestine (middle). For WT virus infection, small intestinal lamina propria blood vessel congestion, infiltration and edema resulting in swelling of the villi (solid arrows) and enterocyte desquamation (open arrows) were observed.

**Figure S3.**
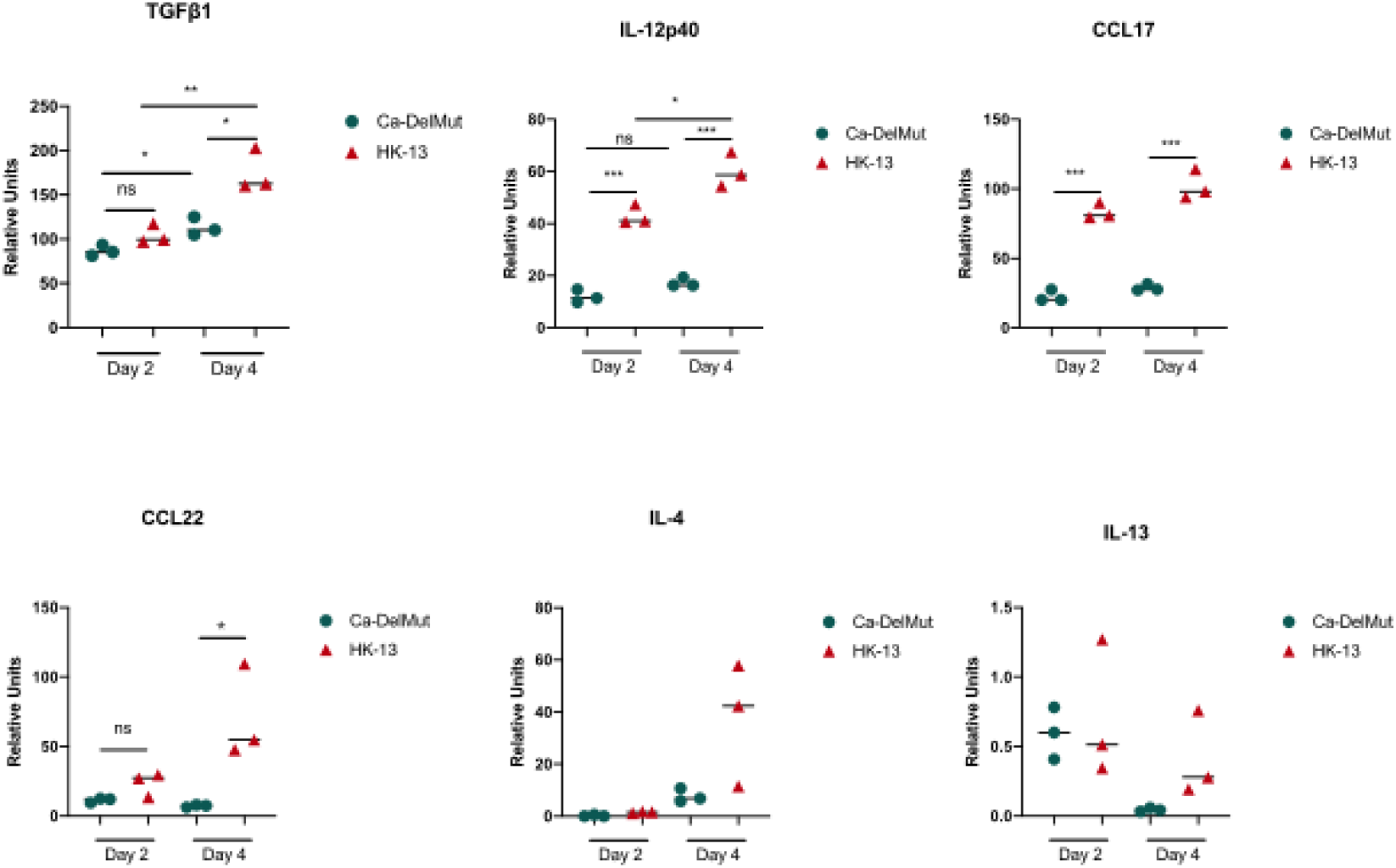
Ca-DelMut does not induce elevated levels of proinflammatory cytokines in hamsters. Hamsters were infected intranasally with 1×10^3^ pfu of either Ca-DelMut or WT HK-13 virus. At days 2 and 4, RNA was extracted from lung and nasal turbinate tissues using RNAsol RT procedures. cDNA was synthesized using oligo dT primers. Expression of different proinflammatory cytokines was examined by qPCR, normalized to the internal reference gene (hamster γ-actin), and the comparative Ct (2-ΔΔCt) method was utilized to calculate the cytokine expression profile. Statistical comparisons between means were performed by Student’s t-test: *** p<0.001, ** p<0.01, * p<0.05, ns: not significant.

**Figure S4.**
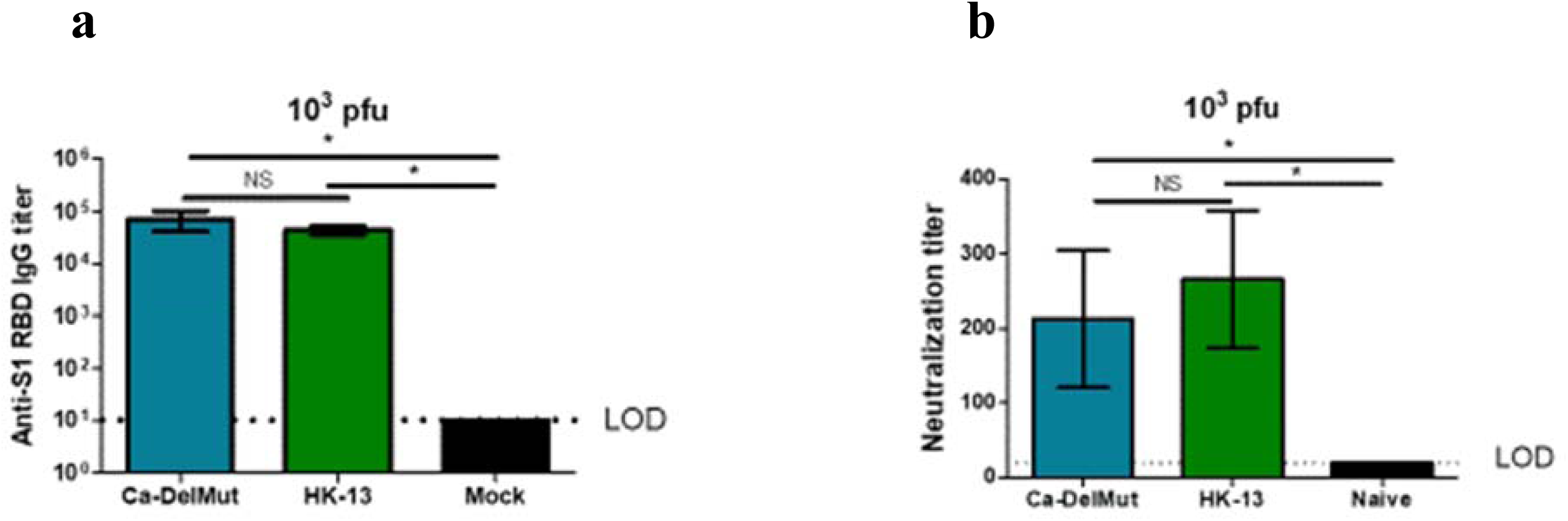
Lower dose (10^3^ pfu) Ca-DelMut immunization is able to induce a strong humoral response in hamsters. Hamsters were infected intranasally with 1×10^3^ pfu of either Ca-DelMut or HK-13 virus, or mock immunized. At day 21, blood was collected from hamsters and tested for anti-S1 RBD IgG titers (a) and neutralization activity against HK-13 virus (b). Error bars represent mean ± s.d. (n=3). LOD: level of detection.

**Figure S5.**
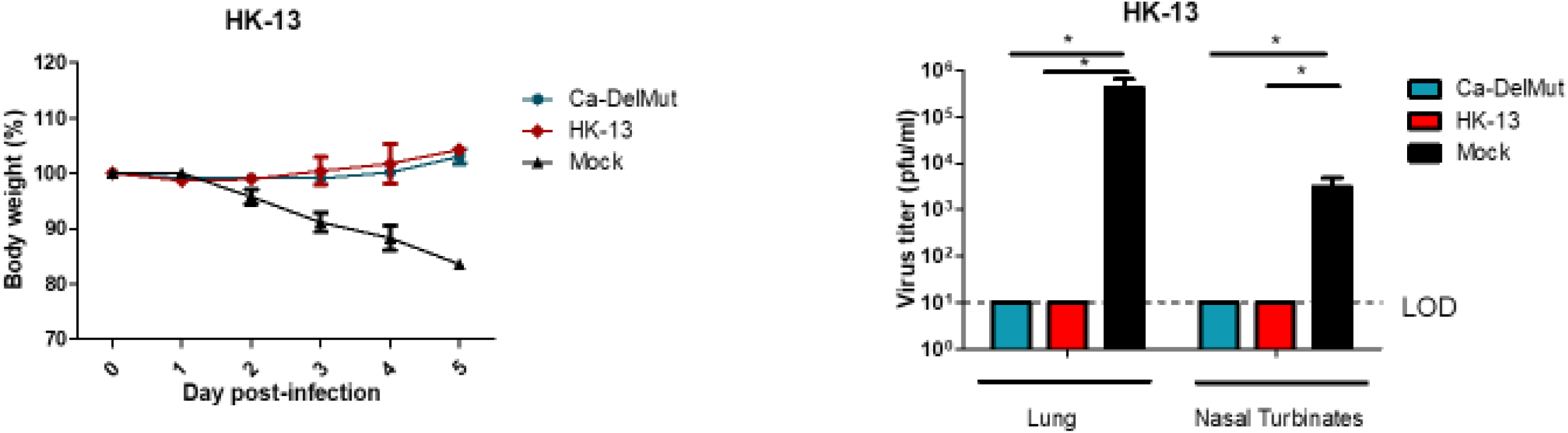
Lower dose Ca-DelMut immunization still provides complete protection against WT HK-13 virus. Hamsters were infected intranasally with 1×10^3^ pfu of either Ca-DelMut or HK-13 virus, or mock immunized. At day 28 after immunization, hamsters were challenged with 1×10^3^ pfu of HK-13 virus. Body weight change and disease symptoms were monitored for 5 days. At day 5 post-infection, lungs and nasal turbinate tissues were collected for virus titration. Error bars represent mean ± s.d. (n=3).

**Figure S6.**
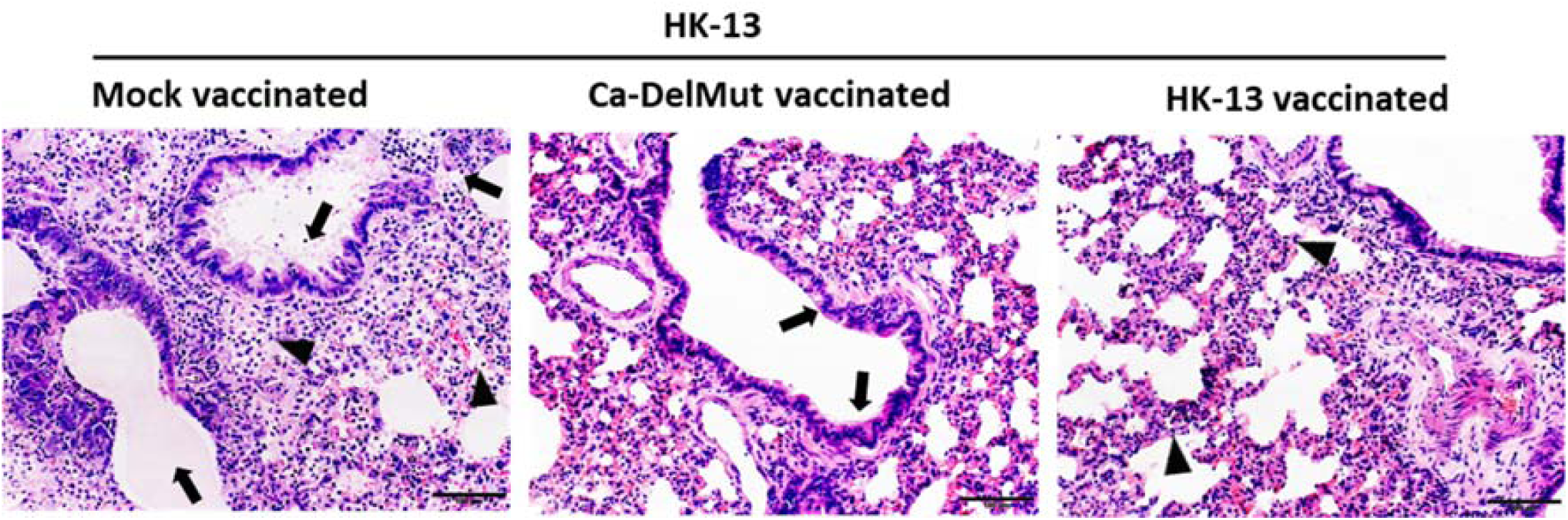
Low inoculum of Ca-DelMut provides protection againstreinfection of wild type SARS-CoV-2. Hamsters were infected intranasally with 1×10^3^ pfu of either Ca-DelMut, HK-13 strain or mock immunized. At day 28 after immunization, hamsters were challenged with 1×10^3^ pfu of HK-13 virus. At day 5 post infection, lungs were collected for histopathological study. Lungs were fixed in 10% formalin, and then processed in paraffin blocks and H&E staining. Mock vaccinated hamster lung showed bronchiolar epithelial cells death and luminal secretion mixed with cell debris (arrows); diffuse alveolar infiltration and exudation (arrowheads); Ca-DelMut immunized lung showed no apparent bronchiolar epithelium cell death (arrows), alveoli showed regional septal infiltrationIn HK-13 strain immunized hamster lung showed no apparent histopathology other than alveolar wall thickening (arrowheads) after re-challenge.

